# The Host Fruit Amplifies Mutualistic Interactions between *Ceratitis Capitata* Larvae and Associated Bacteria

**DOI:** 10.1101/327668

**Authors:** Doron Shalom Yishai Zaada, Michael Ben-Yosef, Boaz Yuval, Edouard Jurkevitch

## Abstract

Background:

The Mediterranean fruit fly *Ceratitis capitata* is a major pest in horticulture. The development of fly larvae is mediated by bacterial decay in the fruit tissue. Despite the importance of bacteria on larval development, very little is known about the interaction between bacteria and larvae in their true ecological context. Understanding their relationship and inter-dependence in the host fruit is important for the development of new pest control interfaces to deal with this pest.

Results:

We find no negative effects on egg hatch or larval development brought about by the bacterial isolates tested. The various symbionts inhabiting the fly’s digestive system differ in their degree of contribution to the development of fly larvae depending on the given host and their sensitivity to induced inhibition caused by female produced antimicrobial peptides. These differences were observed not only at the genus or species level but also between isolates of the same species. We demonstrate how the microbiota from the mother’s gut supports the development of larvae in the fruit host and show that larvae play a major role in spreading the bacterial contagion in the infected fruit itself. In addition, we present (for the first time) evidence for horizontal transfer of bacteria between larvae of different maternal origin that develop together in the same fruit.

Conclusions:

Larvae play a major role in the spread and shaping of the microbial population in the fruit. The transfer of bacteria between different individuals developing in the same fruit suggests that the infested fruit serves as a microbial hub for the amplification and spread of bacterial strains between individuals.

## Background

According to the hologenome theory, multicellular organisms and their associated microorganisms form individual holobionts in which the host and its symbionts act as a consortium; the ability of the microbiota to rapidly adapt to novel conditions endows the combined holobiont with greater adaptive potential than the one provided by the host’s own genome[1].

In insects, bacterial associations are ubiquitous and indubitably have contributed to the impressive success of this group, which dominates terrestrial ecosystems [2–4].

Symbiotic microorganisms have been implicated in several critical processes that increase the fitness of their insect hosts (reviews by [5–7]). Most important among these functions is nutrition, whereby primary, obligate symbionts provide hosts with otherwise unavailable nutrients. Furthermore, secondary, facultative symbionts, which may also provide essential nutrients to their hosts, contribute to a wide array of beneficial traits, such as adaptation to thermal stress, resistance to pathogens, insecticides, predators and natural enemies (e.g. [7–11]), dispersal and increase in host range [12,13]. In addition to providing models for examining explicit evolutionary and functional hypotheses, these symbioses can be manipulated in efforts to control vectors of disease and economically important pests (reviews by [14–17]).

True fruit flies (Diptera: Tephritidae) develop in the tissues of host plants, particularly ripening fruit. A key event in the evolution of this group of flies was the departure from saprophagy (feeding on decaying, spoiled tissues) to feeding on live plant tissue [18].

The brokers of this switch (*sensu* Douglas [19]), which opened up a new adaptive landscape for the flies, were rot inducing bacteria that established successfully in the living tissue of the plant (discussed by Ben-Yosef et al. [20,21]). The developing fruit presents a nutritionally challenging environment, low in protein yet high in sugar, as well as myriad secondary metabolites and structural challenges whose goal is to deter phytophages. Gut bacteria of fruit flies, maternally transmitted during oviposition, have been implicated in the development of larvae in fruit, either through overcoming plant defenses [21] or through pectinolytic and diazotrophic activities that compensate for nutritional deficiencies [22].

The Mediterranean fruit fly, *Ceratitis capitata*, a multivoltine and polyphagous species, is one of the most notorious members of the tephritid family, posing a threat to agriculture in many areas of the globe. The gut of this fly hosts a varied yet stable community of bacteria, comprised mainly of several species of the Enterobacteriacae. Species belonging to *Klebsiella*, *Pantoea*, *Enterobacter*, *Citrobacter*, *Pectobacterium* and *Providencia* are commonly found, and have been shown to contribute to pectinolysis in larvae, and in adults, nitrogen fixation, protection from pathogens, and reproductive success (reviewed by Behar et al. [23]).

When female medflies oviposit, eggs are coated with antimicrobial peptides (AMPs) produced in the female accessory gland [24]. Concurrently, the oviposition site is inoculated with bacteria originating in the female gut [22]. This raises two important questions: First- are some members of the bacterial community inimical to egg hatching and subsequent larval development? Secondly, do the AMPs produced by the female selectively favor some bacterial species over others?

Adult fruit flies are winged and highly mobile, and frequently feed on the surface of fruits and leaves, regurgitating gut contents as they do so [25]. Hence it stands to reason that they actively disperse members of the microbiota in the environment (and acquire new ones). The role of larvae in amplifying bacterial populations through their mobility and feeding activity within fruit has not been studied.

The vertical transmission of symbionts, from parents to offspring is common in the insects [26], and has been documented for fruit flies [27]. Horizontal transmission, which has been studied extensively in some hemipterans [13,28,29] has been recently demonstrated (in artificial conditions) for the oriental fruit fly, *Bactrocera dorsalis* [30]. It is very common for numerous medfly females to oviposit, simultaneously or in sequence, in the same host fruit. Thus multiple larvae, originating from different parents, develop within the same fruit. This pattern offers the opportunity for bacteria originating in one parent, to transfer, mediated by decomposing fruit tissue, to unrelated larvae, and subsequently disperse onwards as adults.

In this study, we show that individual bacterial strains isolated from the medfly, some belonging to the same species, differentially affect larval development, experience different sensitivities to egg-antimicrobial compounds, and may be transferred horizontally between con-specific larvae in the fruit.

## Materials and Methods

### Source of bacteria, isolation and identification

We used the previously described N8 streptomycin resistant strain of *Klebsiella oxytoca*, originally isolated from the gut of a wild fly [31,32]. All other bacteria used herein were isolated from the gut of wild females trapped in the vicinity of Rehovot, Israel. Trapped flies were externally sterilized prior to dissection of the gut as previously described [20]. Following dissection, the gut was homogenized and directly plated on diagnostic Chromagar plates (HY Labs, Rehovot). Resulting bacterial colonies having different morphologies and color were isolated, and stocked in 25% glycerol solution at −80°C. Isolates were subsequently identified by sequencing approximately 566bp of the V3 ‒ V5 region of bacterial 16S rDNA (341F-907R primer-pair, *E. coli numbering*) [33]. Sequence similarities were tested against the NCBI (http://www.ncbi.nlm.nih.gov) and SILVA databases (http://www.arb-silva.de) using the Basic Local Alignment Search Tool (BLAST), and SILVA Incremental Aligner (SINA), respectively.

### Effect of bacterial isolate on egg hatch

Freshly laid eggs of ‘Sadeh’ strain Mediterranean fruit flies were obtained from the fruit fly rearing facility of the Israeli Citrus Board. Eggs were surface sterilized in 300 ppm sodium hypochlorite solution, for 2 minutes, followed by double rinsing in 1ml of sterile 0.1M phosphate buffered saline (PBS, pH 6.8). Surface sterilized eggs, were incubated for 10 min in 1ml of PBS containing a single bacterial isolate, or an equal mixture of all examined bacteria adjusted to a density of ~1 O.D (measured at 600 nm). Triplicates of approximately 25 eggs from each treatment group, including control groups of non-treated and surface sterilized eggs were transferred to sterile petri dishes containing sterile solidified agar. Plates were sealed with parafilm and incubated at 27°C for 2 days during which egg hatch was monitored using a stereomicroscope (SteREO Discovery V8; Carl Zeiss MicroImaging GmbH, München, Germany) at 12-hour intervals.

### Effect of antimicrobial peptides on bacterial isolates

Extraction of anti-microbial peptides (AMPs) coating the egg surface was achieved according to previously published protocols [24,34]. Briefly, 250 mg of freshly laid eggs were agitated in 1ml of 0.1 M PBS for 5 minutes, after which eggs were removed by centrifugation. The remaining supernatant was boiled for 10 min and subsequently centrifuged at 10,000g for 10 min to remove proteins of high-molecular weight. The amount of protein remaining in the supernatant was determined using the Bradford protein assay [35] and subsequently adjusted to 100 ng.ml^−1^ by dilution in PBS. The resulting AMP solution was stored at 4°C for up to 48 hours before use.

The effect of AMP extract on bacterial growth was examined by the agar well diffusion method [36]. LB agar plates containing 20 ml of medium (1.7% agar) were seeded with 50μl of bacterial culture (10^6^ CFU.ml^−1^) Using a sterile cork borer, six 5mm-diameter wells were bored into the agar. Subsequently, 50μl of the tested antimicrobial agents were transferred to each well: Two wells contained AMP solution at 100ng protein.ml^−1^, another pair of wells contained AMP solution at 50ng protein.ml^−1^, one well contained 1 mg.ml^−1^ of streptomycin (Sigma) solution in PBS and the sixth well served as a control containing 50μl of sterile PBS. Plates were later sealed and incubated overnight at 27°C. On the following day plates were digitally recorded, and the diameter of the growth inhibition zone surrounding each well was digitally determined using Image J [37]. The response of each isolate to antimicrobial agents was tested on two separate plates.

### Larval contribution to bacterial dispersal

The contribution of larvae to the distribution of bacteria was examined by allowing neonate larvae to disperse on solid LB agar and subsequently monitoring the coverage achieved by bacterial growth on the plate. One, two or three freshly laid eggs of the ’Sadeh’ strain were incubated on sterile solid LB medium, at 27°C for six days, during which hatched larvae were able to freely move throughout the plate. Plates where digitally recorded twice on a daily basis, and the area covered by bacterial colonies was determined by analyzing the photos using ImageJ software. Control plates included 1, 2 and 3 non vital eggs, which were frozen for 4hrs at −20 °C, or eggs which were surface sterilized as described above. Experiments included four replicates for each treatment group, and one replicate for each of the control treatments.

In order to determine whether the number of bacteria in the fruit tissue is correlated with larval development, we used ripe apricot fruits (n= 20). After external disinfection fruits were covered with sterile plastic containers and two V8 female flies were introduced into the containers allowing them to oviposit. Fruit were subsequently maintained at 23°C for eight days, after which larvae were extracted from the fruit, counted and measured for body length under a stereoscope. Additionally, about 300 mg of each fruit pulp were sampled, weighed and homogenized in 1 ml of sterile PBS. Homogenates underwent a series of decimal dilutions in PBS and plated in triplicates on LB agar. Plates were incubated at 37°C for 24 hours and the resulting colonies were counted.

### Effect of bacteria on larval development in fruit

Surface sterilized ‘Sadeh’ strain eggs, were inoculated with each of the 8 examined bacterial isolates or a mix of all isolates by incubation in a suspension of the bacteria, as previously described. Following incubation 30μl of bacterial suspension, containing approximately 15 eggs were injected, under sterile conditions, into a 2 mm deep pore, created with a sterile syringe needle in a surface sterilized, fresh plum (*Prunus salicina*) fruit. Each fruit was pierced and injected twice: once in each side. Each isolate and the mixed suspension of all bacteria were tested in two fruits (four injections total). Control fruit (n=3, Six injections total) were inoculated with sterile PBS containing surface sterilized eggs. To prevent egg desiccation, pores where sealed with 10μl of 2% sterile agar immediately after injection. The infested fruit were incubated for eight days in a sterile laminar flow cabinet at room temperature. Subsequently, fruits were dissected using a sterile blade and all larvae were extracted, counted and measured. The contribution of bacteria to larval development was determined by comparing average larval length between each of the treatments and the control group.

### Fruit mediated horizontal transfer of bacteria

Three ripe surface sterilized peach fruits (*Prunus persica*) were exposed to simultaneous oviposition by wild females fed on streptomycin resistant strain of *K. oxytoca* (N8) (N8W) and axenic mass-reared Vienna 8 (AxV) females. The axenic (bacteria free) condition was achieved as described by Ben-Yosef et al.[38] A fourth fruit was exposed only to oviposition by AxV flies, and served as a control. All females mated prior to the beginning of experiments. Larvae were extracted from fruit five days after oviposition, surface sterilized with 70% ethanol, and aseptically dissected to extract the gut. Individual whole guts were homogenized in 50 μl sterile PBS and plated both on LB and selective LB (that contained 500microgram.ml^−1^ streptomycin) solid medium plates. Media were incubated for 24 hours at 27°C incubator. Upon successful colonization of gut extraction on selective LB medium we used the dissected larvae to determine its maternal origin. DNA extraction of the larval tissue was performed using DNeasy blood and tissue kit (Qiagen, Hilden Germany) according to manufacturer instructions. DNA was amplified by PCR using the CCmt primer pair (Ccmt5495, AAA TCA CCA CTT TGG ATT TGA AGC; and Ccmt5827, TGA AAA TGG TAA ACG TGA AGA GG) targeting flanking regions of tRNA-Gly of the medfly mitochondrial genome. Amplification product was cut with the HaeIII restriction enzyme (Takara-Bio, Otsu, Japan) targeting a polymorphic distinguishing the WT and the V8 strains (for a detailed description see San Andres et al., [39]). Prior to the experiment the protocol was validated on 50 V8 and wild females (results not shown).

The identity of colonies resistant to streptomycin was determined by sequencing the 16S rRNA (between bp 341 to 907) as previously described.

### Statistical analysis

Parametric tests were applied where datasets were normally and homogenously distributed. Otherwise, non-parametric tests (Wilcoxon signed-rank test) were used.

Tukey-HSD and ANOVA tests where used to establish differences in the response of hatching ratio to bacteria, AMP on bacteria and the effect of larvae numbers on the distribution of bacteria. Linear regression was applied to test correlations between number of larvae and larval length or bacteria titer in fruit tissue.

Statistical significance was set at ◻= 0.05, but when multiple comparisons were needed Bonferroni correction was applied.

Data processing and analysis was performed using JMP pro v. 10 statistical package (SAS, Cary,NC, USA). Means and their co-responding standard errors are reported.

## Results

### Effect of bacterial isolate on egg hatch

To examine the effect of bacteria on egg viability, eggs which had been exposed to different bacterial isolates were incubated for 48 hours, after which hatching ratio was recorded for each treatment. Following incubation 83.4% of all eggs had hatched and no further eclosions were observed. Treatment had a significant effect on egg hatch ratio (ANOVA, F_12,38_=4.256, *P*=0.001; Tukey’s HSD, *P*<0.05, figure 1). Untreated eggs (UT) had the lowest hatching rate (60.9%) which significantly differed from all other treatment groups, excluding eggs which had been exposed to a mixed bacterial culture (Mix) (Tukey’s HSD, *P* <0.043, *P* >0.055 respectively). These eggs eclosed at a higher rate (80.5%) but remained statistically inseparable from all other groups (Tukey’s HSD, *P* >0.0695, Figure 1). Eggs which had been exposed to single bacterial isolates were not affected by the type of bacteria (ANOVA, F_9,20_=0.924, *P*>0.525; Tukey’s HSD, *P* >0.618), and had a similar hatching rate to that of surface sterilized eggs (SHC treatment, 81.8% to 93.3%, Tukey’s HSD, *P* >0.766). Eggs incubated with *Citrobacter freundii* III and *Pseudomonas aeruginosa* bacteria had a relatively low hatching rate (81.8% and 82.5% respectively, figure 1), while the highest hatch ratio was for eggs exposed to *Citrobacter werkamnii* (93.32%, figure 1).

**Figure 1.**
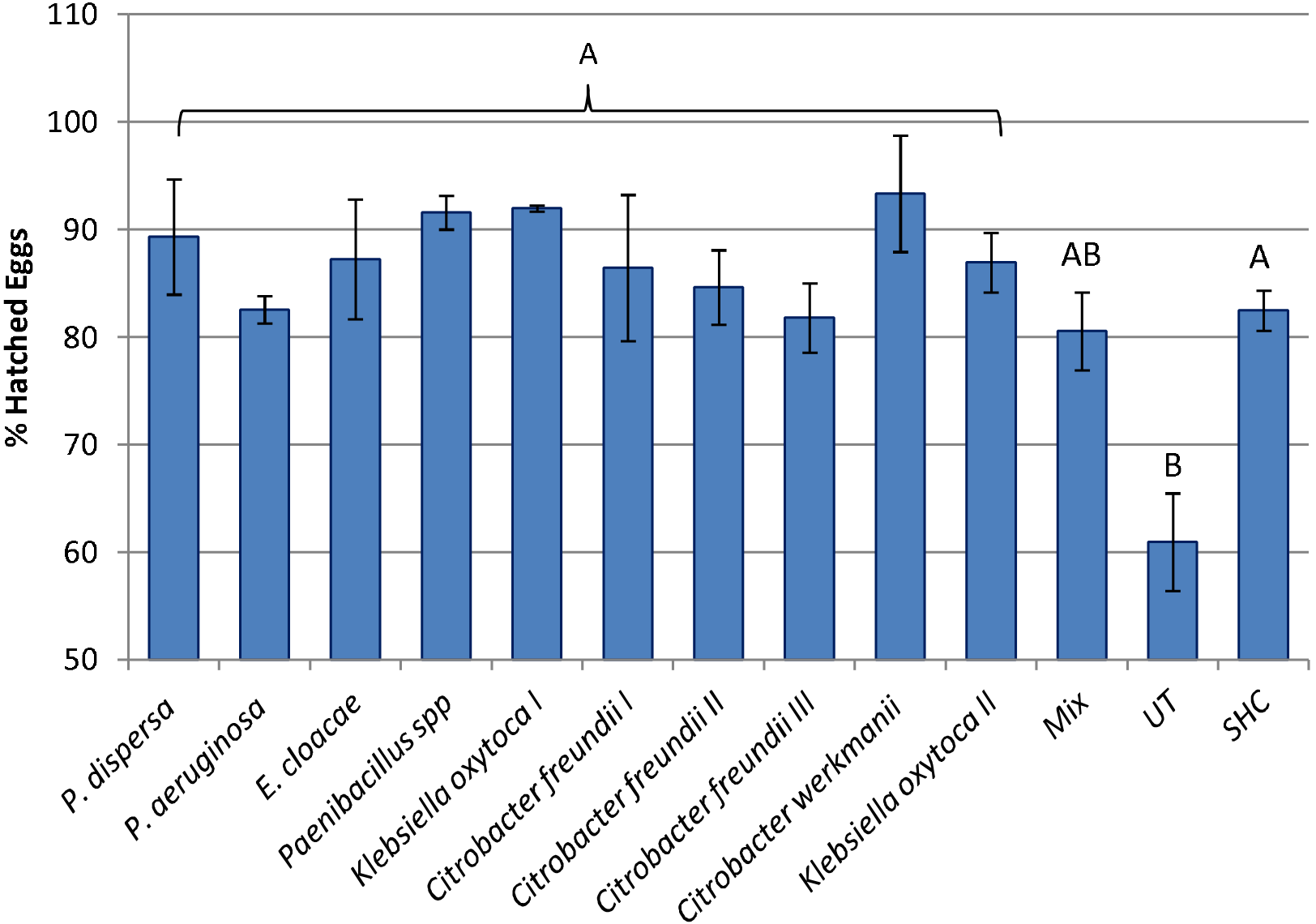
Effect of bacterial strain on egg hatch. Percentage of eggs hatching when inoculated by single or mixed (mix) bacterial strains isolated from the medfly, eggs treated with sodium hypochloride (SHC) or untreated (UT). Means denoted by different letters are statistically different (Tukey’s HSD *P*<0.05)

### Effect of antimicrobial peptides on bacteria

Extracts containing AMPs inflicted an inhibitory effect to the vast majority of the challenged isolates (10 out of 11). Similarly, streptomycin inhibited the growth of ten of the tested isolates, excluding one isolate (*Paenibacillus* sp.), that was unaffected by the antibiotic. The inhibition zone around streptomycin wells was consistently larger (16.68±0.62mm) than those surrounding wells filled with AMPs solution (5.14±0.3mm) (T_98_=20.44, P<0.0001). There was no difference in halo size between 50 mg.ml^−1^ (4.95±0.43) and 100 mg.ml^−1^ (5.32±0.42) (T_77_.9=0.65, P=0.54). While most isolates were inhibited to some extent by the antimicrobial agents, some exhibited a remarkable response. *Paenibacillus* sp., the single isolate not to be affected by streptomycin, demonstrated the highest susceptibility to AMPs (inhibition halo diameter >10mm), *Pseudomonas aeruginosa* was the only isolate that was unaffected by the application of AMPs. None of the control sites, containing PBS, exhibited any inhibition.

With the exception of the two extremes, the tested isolates exhibited a variety of responses to the AMPs, which was evident both at the species and strain levels. Thus, the lowest sensitivity was found in 2 of the *Citrobacter freundii* isolates tested, while the highest sensitivity was found in the third strain of this species (Figure 2). In another case, 2 strains of *K. oxytoca* were inhibited uniformly by streptomycin, yet differed in their response to AMPs (Figure 2).

**Figure 2.**
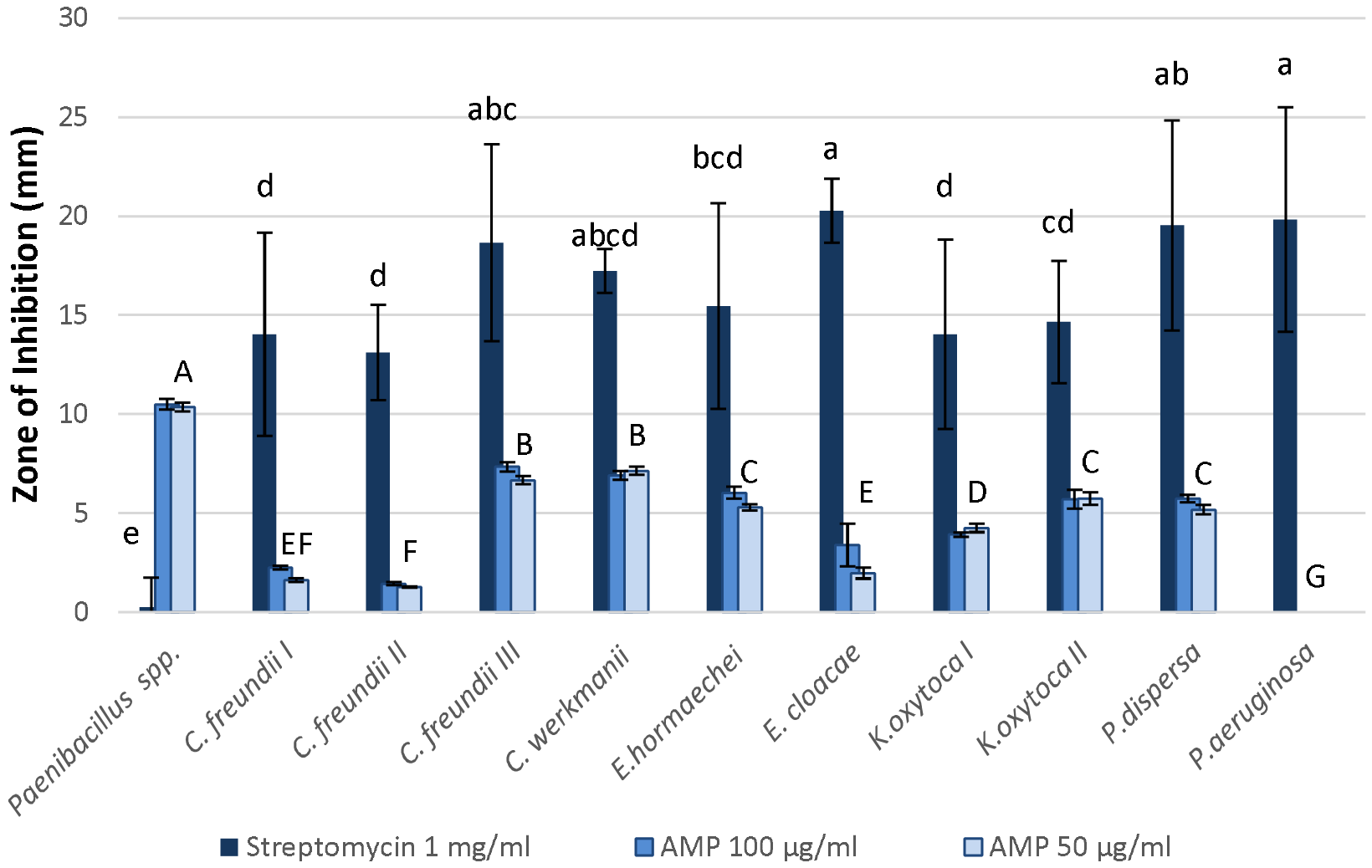
Suppressive effects of antimicrobial peptides extracted from medfly eggs (AMP) and antibiotics (streptomycin) on intestinal bacterial strains. Intensity of antimicrobial activity is measured as the diameter of the bacteria-free zone surrounding wells containing 50 μl of the examined solution. Columns denoted by different letters are statistically different (Tukey’s HSD *P*<0.05). Comparisons of the response to AMP and streptomycin are indicated by capital or lower case letters, respectively. The response to AMP was independent of the concentration and thus represented by a single letter for both columns.

### Larval contribution to bacterial dispersal

The wandering of larvae on a growth medium brought about bacterial dispersal. Increase in the number of larvae resulted in increased bacterial dispersal, measured as the percentage of the plate covered by bacterial growth. This was highest (41.26 ± 0.78%) in the treatment containing two larvae. This percentage significantly differed from the plates that contained three larvae (27.31±3.25%) and one larva (20.63±1.62%) (Tukey HSD P<0.001). In the first eight hours of the experiment, microscopic colonies were observed in proximity to the egg placement area of each treatment group, at this stage no larvae were observed. After 21 hours, except for the freeze treatment, all eggs were hatched, yet spread of the bacterial inoculum was observed in only one of the plates, in the treatment containing 3 eggs. Starting with the fourth observation (41 h post placement), evidence of bacterial dispersal was confirmed in all treatments, and the percentage of colony coverage increased steadily throughout the experiment. In the fifth observation (56 h), the average coverage area of the plates containing 2 and 3 larvae was over 15%, whereas in the parallel treatment containing single larvae, less than 4% coverage was recorded. However, a difference in the area covered between the various treatments was recorded only in the sixth observation (62 h), where the percentage of coverage of the plates in which 2 larvae roamed differed from those containing a single larva. From this point on, throughout the experiment, the differences between the plates containing two larvae and those containing one were preserved, and in the last two observations, the first was distinguished (Tukey HSD P<0.001) from the treatment containing 3 larvae (Figure 3). At no stage was bacterial growth or spread observed in any of the control treatments.

**Figure 3.**
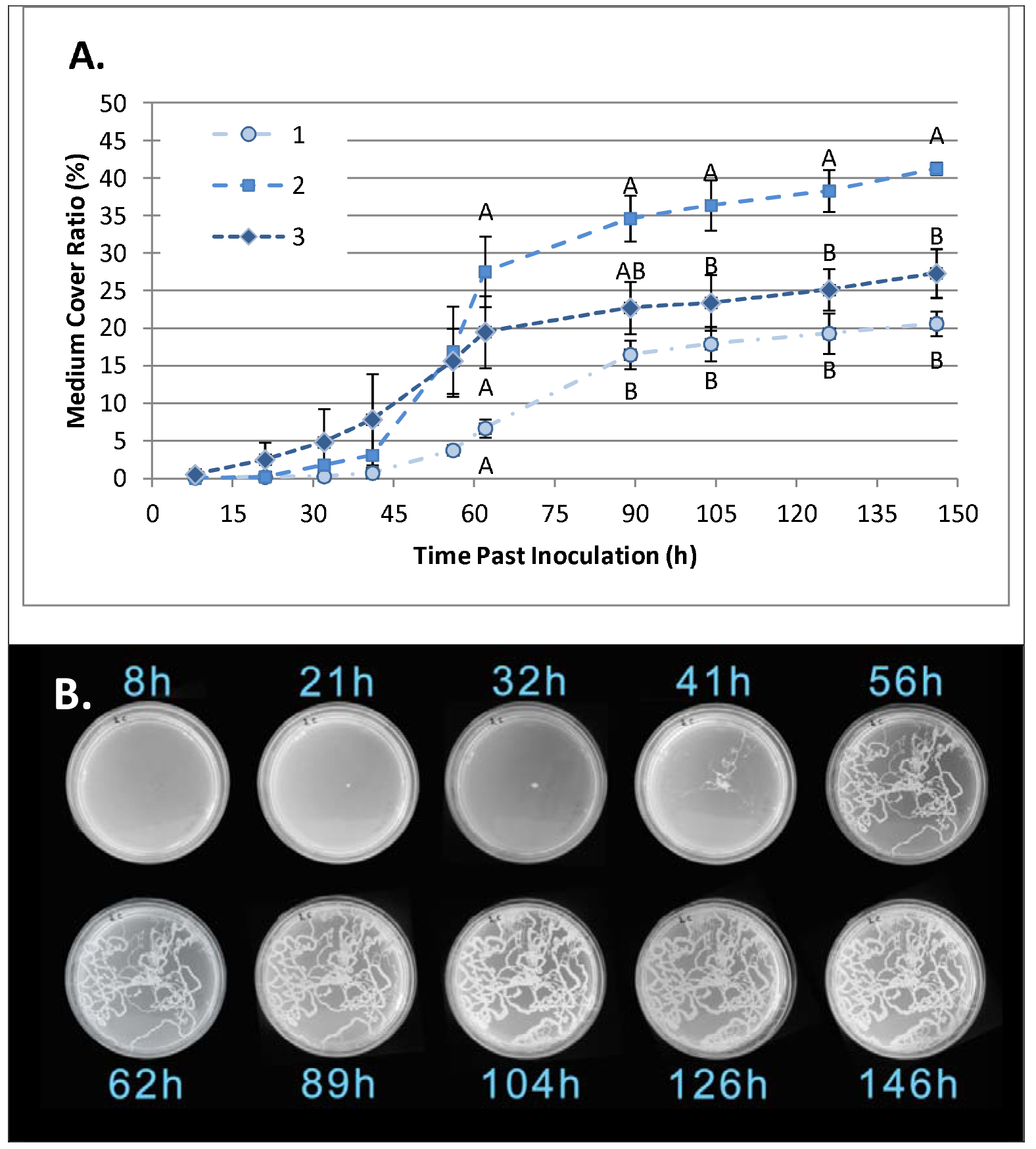
Larvae-mediated dispersal of bacteria. A. Bacterial growth, measured as a function of time (as % of total surface) following the placement of one, two or three medfly eggs onto a Petri dish containing solid LB is presented as % of total surface area. Differences between groups were established separately for each time point. Different letters denote significant differences between groups for each time point (Tukey’s HSD *P*<0.05). B. Time-lapse photographs of a single plate containing two larvae. The spread of bacteria is clearly visible by trails of developing colonies depicting the movements of advancing larvae.

A similar pattern emerged *in vivo:* In apricot fruits, the number of bacteria correlated with the number of developing larvae. The number of larvae in the fruit ranged from 2 to 73 (average 35.93 ± 6.15) and the amount of bacteria in the tissue of the fruit ranged from 1396 to 2.4 • 10^8^ CFU. g^−1^ (Figure 4). There was a significant logarithmic correlation between total larvae in fruit and the CFU. g^−1^ (R^2^ = 0.46, F8 = 5.97, P = 0.044). No correlation was found between the logarithm or the number of colonies per gram fruit and larval length (R_2_ = 0.01, F_8_ = 0.05 P = 0.819), nor to the number of larvae and their length (R_2_ = 0.13, F_8_ = 1.06, P = 0.336). These results are based on data obtained from 20 fruits that contained a total of 528 larvae (Figure 4).

**Figure 4.**
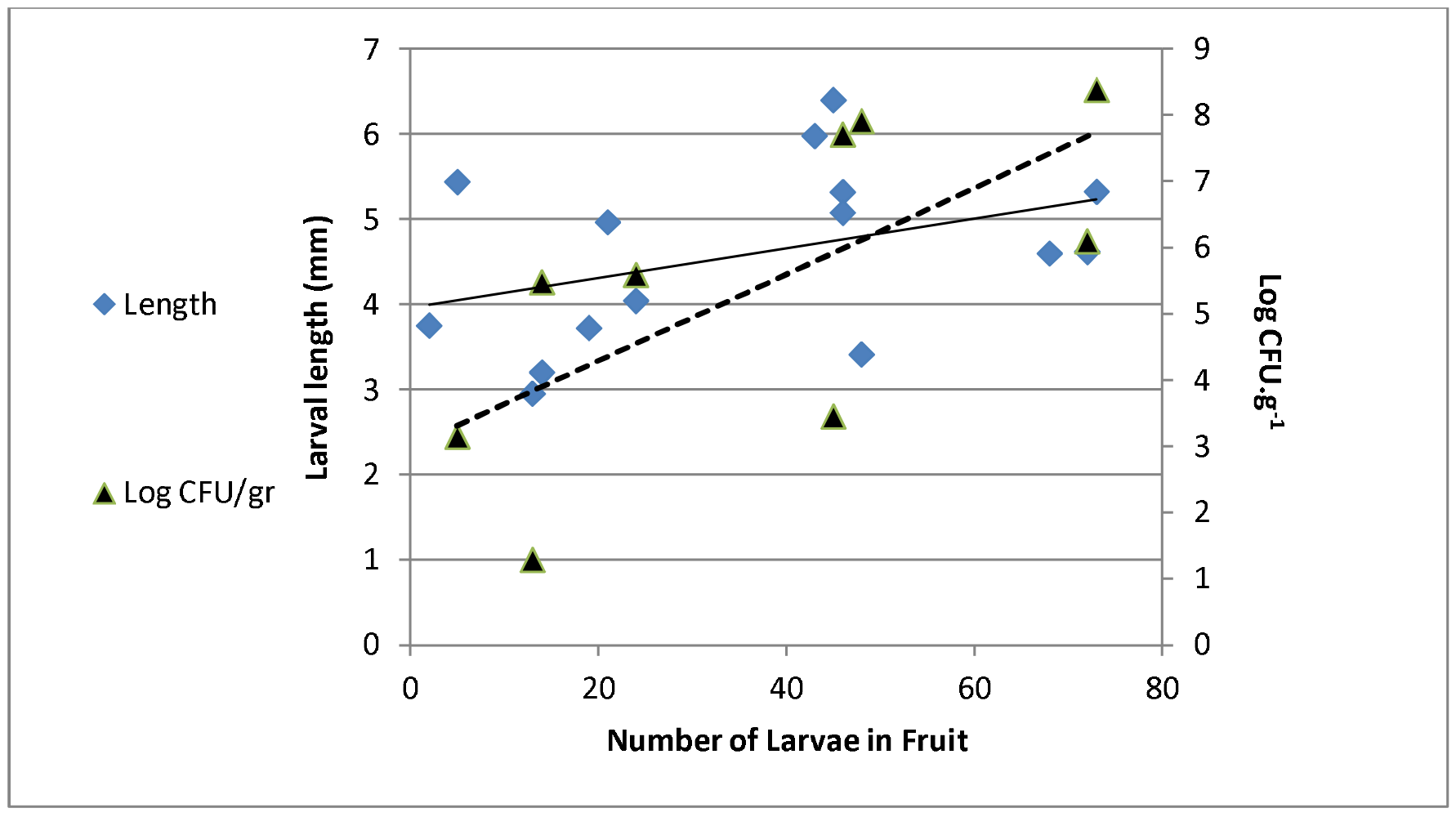
Effect of larvae on bacterial abundance in fruit. Average larval length (blue rectangles) and concentration of bacteria (as CFU.g^−1^Fruit pulp) (black triangles) as affected by the number of larvae developing in apricot fruits. The bacterial titer was significantly correlated with the number of larvae in fruits (*P* = 0.044). Larval length was not significantly correlated with the number of larvae developing in the fruit (*P*=0.336).

### Effect of bacteria on larval development in fruit

Different isolates resulted in different effects on larval length. Some of the isolates had a positive effect on larval length, in comparison to the control treatment, and no negative effect was observed (Figure 5). Isolate identity did not affect the number of vital larvae extracted from fruits at the end of the incubation period (ANOVA F_9,16_=0.72 P=0.685), but had a significant effect on larval length (Welch’s F_9_=36.45 P<0.0001).

**Figure 5.**
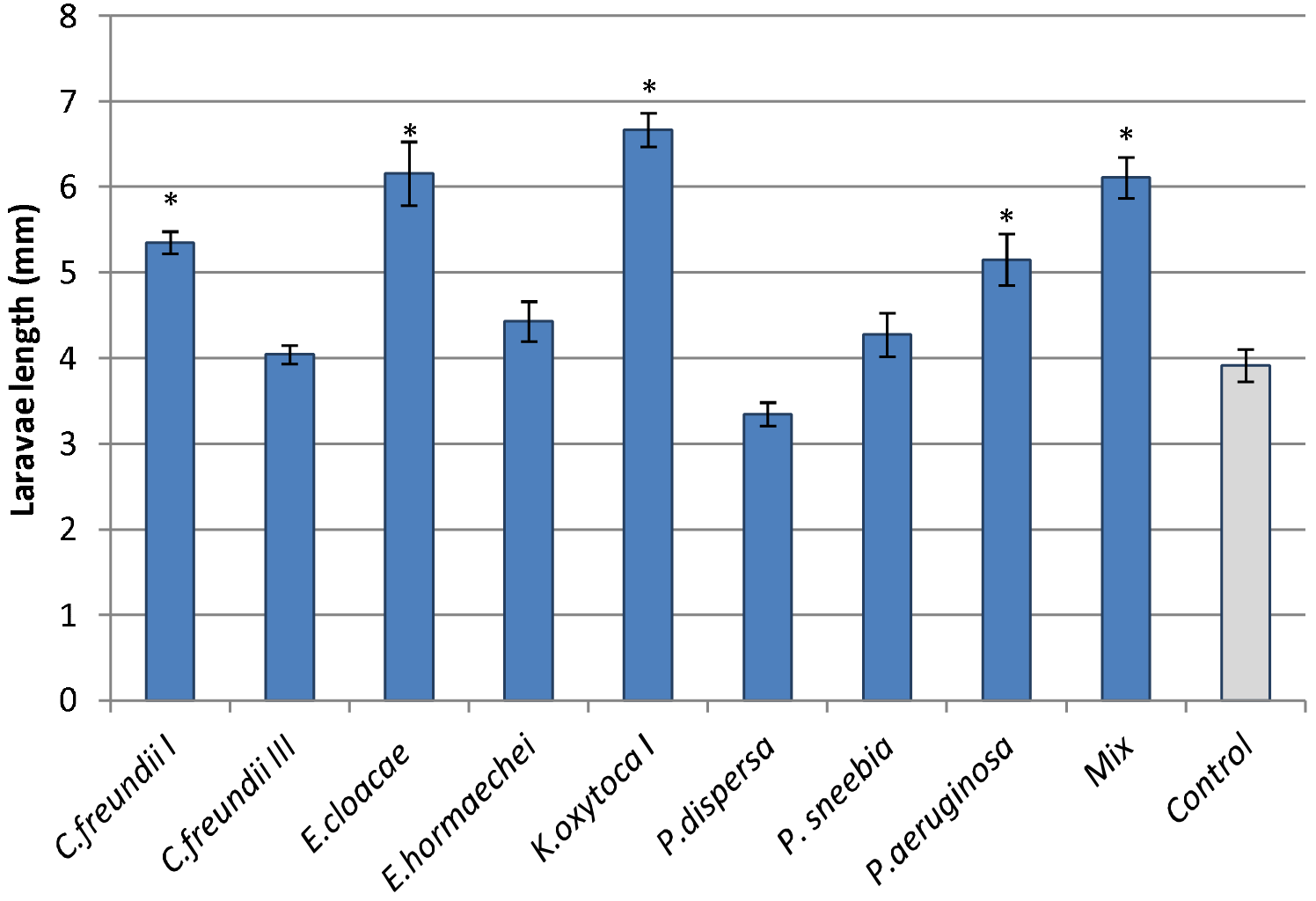
Effect of bacterial strains isolated from the medfly on the average length of larvae developing in fruit. Surface sterilized eggs incubated in a pure culture of each isolate or in an equal mixture of all isolates (Mix), all in PBS, were subsequently inoculated into plums. Larval length was recorded after eight days. Control eggs were treated with sterile PBS Treatments differing significantly from the control are denoted by asterisks (Wilcoxon signed rank test, Z= −4.23, P<0.0055).

Of the eight isolates tested, four significantly contributed to larval development (in terms of body length) compared to the aseptic control treatment (3.916±0.187) (Wilcoxon signed rank test, Z= −4.23, P<0.0055). The largest larvae derived from fruits infested with eggs inoculated with *K. oxytoca* (6.66± 0.16), and *E. cloacae* (6.15 ±0.3). Eggs inoculated with *Pantoea dispersa* and *Citrobacter freundii* III resulted in the lowest larval development rate, reaching 3.34±0.13mm and 4.04±0.11 mm respectively, and did not differ from the aseptic control (Wilcoxon signed raneked test Z> −1.96 *P*>0.049). Larvae developed from eggs incubated with the microbial mixture reached an average length of 6.11±0.25mm and differed significantly from the control (Wilcoxon signed rank test, Z= −5.44, P<0.0001).

### Fruit mediated horizontal transfer of bacteria

In this experiment, peach fruits were exposed to simultaneous oviposition by wild female flies fed on a diet enriched with an antibiotic resistant bacteria strain, and an axenic V8 fly. With the exception of one larva, bacteria were detected in all larval gut extracts plated on LB (n=43). The growth of colonies on streptomycin-containing LB was less common (n=16). In each of the three experimental fruits that were exposed to simultaneous oviposition, we found that larval offspring of the V8 axenic females were associated with bacteria which developed on selective media, indicating the acquisition of antibiotic-resistant bacteria from the WT con-specifics. In the control fruit, which were exposed to axenic females only, none of the developing larva were associated with streptomycin-resistant bacteria (Figure 6).

**Figure 6.**
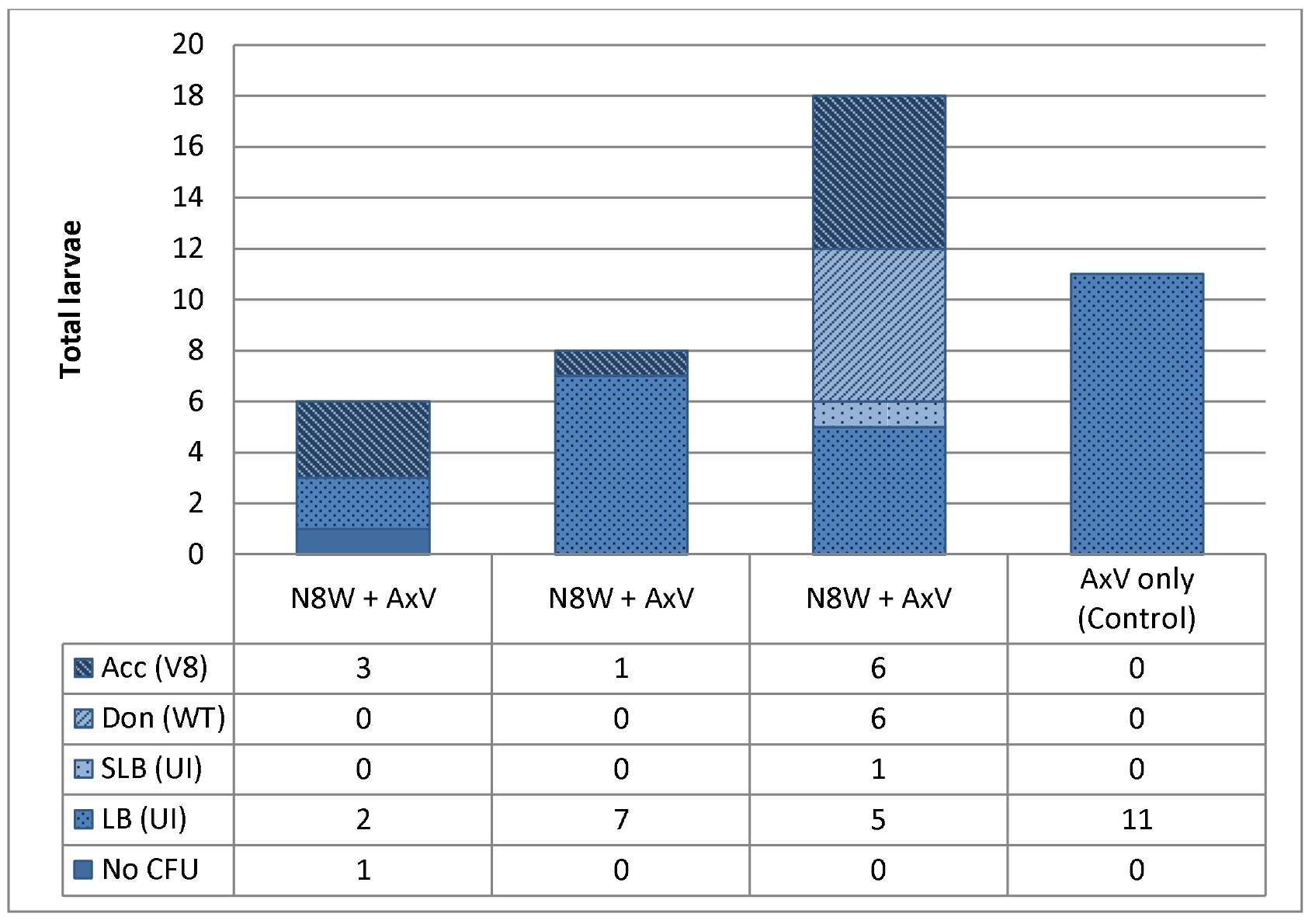
Fruit-mediated transfer of bacteria between conspecific larvae. Transfer of streptomycin-resistant *Klebsiella oxytoca* N8 between WT, field-caught donor flies (N8W) and axenic strain V8 acceptor flies (AxV). The donor and the acceptor oviposited in the same fruit. Larval gut homogenates were plated on selective and non-selective LB media plates. Larvae whose homogenate established on selective media were genotyped. Each column represents a fruit and all the larvae extracted from it, and is designated by the maternal oviposition types (N8W, AxV). Columns are divided according to the various larval genotypes and microbial phenotypes identified. Acc (V8): progeny of AxV mothers, bearing S resistant bacteria; Don (WT): Progeny of N8W mothers, bearing S resistant bacteria; SLB (UI): larvae of unidentified genotype, bearing S resistant bacteria; LB (UI): Larvae with only non S resistant bacteria; No CFU: larvae that yielded no bacterial colonies on either medium.

## Discussion

Drew & Lloyd [40] were the first to recognize that the host plant serves as an activity hub for fruit flies and their associated bacteria. Since then quite a large body of research has focused on the effects of the microbiota on adult fly fitness and on larval development [23]. In this study we focused on the interaction between larvae and bacteria within the host fruit, an interaction we perceive as being of crucial ecological importance for all three participants.

The lowest rate of egg hatch was found in untreated, fully symbiotic eggs (Figure 1). While this may seem paradoxical, we must recall that these are mass reared eggs that bear an excessive bacterial load, one that is not typical of the natural microbiota [31]. Inoculating dechorionated eggs with members of the native microbiota, rescued them from this deleterious artifact (Figure 1). The structure of the bacterial community developing in the fruit is primarily determined by the AMPs present on the egg. Indeed, our results demonstrate how the AMPs produced by ovipositing females constrain the microbial community inoculated into the fruit. The newly hatched larva, through its movement and maceration of fruit tissue, becomes the major agent for distributing bacteria in the host. Thus the fruit becomes a temporary active arena that provides for amplification of bacterial communities and their horizontal transfer between insects.

Selective inhibition by AMPs creates a bottleneck for bacterial diversity in the host, by favoring some species and suppressing others. Changes were also observed at the strain level, where bacteria of the same species respond differently to the AMPs. These results confirm previous findings by Marchini et al.,[41], that described different inhibition response of *K. oxytoca*. We find that this selectivity correlates with the contribution (or lack thereof) of the affected bacteria. The isolates which were least affected by the AMPs were also those that contributed most to larval development in fruit (*K. oxytoca I*, *C. freundii I*, *E. cloacae*, *P. aeruginosa*). Conversely, isolates inhibited by AMPs were also those that least contributed to larval development (Figures 2 & 5). No such effect was found on the contribution of these isolates to egg hatching rates.

We find conclusive evidence for horizontal transfer of bacteria within the fruit (Figure 6). This finding extends the observation of Guo et al.,[30] (who demonstrated horizontal transmission between larvae of *B. dorsalis* developing in artificial media), to host fruit, and highlights the importance of the host fruit as a hub for amplifying and dispersing bacterial populations. Indeed, bacteria capable of jumping ship and moving horizontally to a new invertebrate host will have increased probability of survival [42]. Establishment of larvae in the fruit results in progressive fruit rot, whereby bacterial populations are amplified. In this context it is important to recall that oviposition sites, abrasions and wounds attract adult flies seeking food and oviposition sites [25,43]. Thus, the amplification of bacteria within the fruit, compounded by horizontal transfer, allows adult flies to acquire bacterial isolates from decomposing fruit. In the case of the polyphagous and widely dispersed medfly, this mechanism may equip adult females with novel genetic material, providing the holobiome’s offspring with an enhanced capacity to develop in hosts which differ in their nutritional quality and biochemical defenses and to adapt to other biotic and abiotic fluctuations.

Once infested by medfly larvae and associated bacteria, a successional process begins in the fruit, as it becomes available to insects incapable of breaching the defenses of an intact fruit. In fruit infested by medflies we have seen that these consist initially of various Drosophilids and finally Staphylinid beetles (Yuval, unpublished). Thus, a potential biocontrol strategy would be to target the infested fruit by specific entomopathogens delivered by drosophilids, effectively truncating the medfly life cycle. Future work will determine the feasibility of such an approach.

In this study we studied interactions between medfly larvae and bacteria in host fruit. This provides a degree of ecological realism to our results and conclusions. We used three different host plants to demonstrate different aspects (larval development, bacterial dispersal and horizontal transmission) of this interaction. However, we must bear in mind that the reality in the field is far more complex. The fruit we used were bought in a store, they were in an advanced stage of ripening and probably low on defensive compounds. In the field, female medfly encounter host fruit at earlier stages of maturation, when nutrients are relatively low and the concentration of defensive metabolites high. Accordingly, larval survival is lower in such fruit [44,45]. Furthermore, under laboratory conditions, the natural enemies and competitors are absent. Including these factors (nutrition, parasitism, competition) in future experiments will surely broaden our understanding of the intricate web created between fly larvae, the bacteria they arrive with or acquire, and the host fruit.

## Conclusions

Larvae play a major role in the distribution and shaping of the microbial population in the fruit. The transfer of bacteria between different individuals developing in the same fruit suggests that infested fruit serve as a microbial hub for the amplification and distribution of bacterial strains between individuals. Furthermore, such infested fruit emerge as a promising target for controlling the fly population by introduction of entomopathogenic microbes.

## Declarations

Ethics and consent ‒not applicable.

Availability of data and material - The datasets generated and analysed during the current study are available from the corresponding author on request.

Competing interests- The authors declare that they have no competing interests.

Funding- Supported by grants from The IAEA\FAO, and the Israel Science Foundation (736\16), to BY and EJ.

Authors’ contributions ‒ All authors contributed ideas to formulating the hypotheses tested in this study. Experiments were performed by DZ. Results were analyzed primarily by DZ and MBY. All authors contributed to writing the manuscript. All authors read and approved the final manuscript.

